# Saccharomycotina yeasts defy longstanding macroecological patterns

**DOI:** 10.1101/2023.08.29.555417

**Authors:** Kyle T. David, Marie-Claire Harrison, Dana A. Opulente, Abigail L. LaBella, John F. Wolters, Xiaofan Zhou, Xing-Xing Shen, Marizeth Groenewald, Matt Pennell, Chris Todd Hittinger, Antonis Rokas

**Affiliations:** Department of Biological Sciences, Vanderbilt University, Nashville, TN 37235, USA; Evolutionary Studies Initiative, Vanderbilt University, Nashville, TN 37235, USA; Laboratory of Genetics, J. F. Crow Institute for the Study of Evolution, Center for Genomic Science Innovation, DOE Great Lakes Bioenergy Research Center, Wisconsin Energy Institute, University of Wisconsin-Madison, Madison, WI 53726, USA; Department of Biology, Villanova University, Villanova PA 19085, USA; Department of Bioinformatics and Genomics, University of North Carolina at Charlotte, Charlotte NC 28223, USA; Guangdong Laboratory for Lingnan Modern Agriculture, Guangdong Province Key Laboratory of Microbial Signals and Disease Control, Integrative Microbiology Research Center, South China Agricultural University, Guangzhou 510642, China; Key Laboratory of Biology of Crop Pathogens and Insects of Zhejiang Province, Institute of Insect Sciences, Zhejiang University, Hangzhou 310058, China; Westerdijk Fungal Biodiversity Institute, 3584 Utrecht, The Netherlands; Department of Quantitative and Computational Biology and Biological Sciences, University of Southern California, Los Angeles CA 90089, USA

## Abstract

The Saccharomycotina yeasts (“yeasts” hereafter) are a fungal clade of scientific, economic, and medical significance. Yeasts are highly ecologically diverse, found across a broad range of environments in every biome and continent on earth^1^; however, little is known about what rules govern the macroecology of yeast species and their range limits in the wild^2^. Here, we trained machine learning models on 12,221 occurrence records and 96 environmental variables to infer global distribution maps for 186 yeast species (∼15% of described species from 75% of orders) and to test environmental drivers of yeast biogeography and macroecology. We found that predicted yeast diversity hotspots occur in mixed montane forests in temperate climates. Diversity in vegetation type and topography were some of the greatest predictors of yeast species richness, suggesting that microhabitats and environmental clines are key to yeast diversification. We further found that range limits in yeasts are significantly influenced by carbon niche breadth and range overlap with other yeast species, with carbon specialists and species in high diversity environments exhibiting reduced geographic ranges. Finally, yeasts contravene many longstanding macroecological principles, including the latitudinal diversity gradient, temperature-dependent species richness, and latitude-dependent range size (Rapoport’s rule). These results unveil how the environment governs the global diversity and distribution of species in the yeast subphylum. These high-resolution models of yeast species distributions will facilitate the prediction of economically relevant and emerging pathogenic species under current and future climate scenarios.

## Introduction

Saccharomycotina is a fungal subphylum as genetically diverse as plants and animals^3^ that occurs across a broad range of environments and metabolic modalities^4^. Yeasts provide a plethora of crucial ecosystem functions, acting as mutualists, parasites, and decomposers^5^. Some yeasts are used as biological pest control while others are pathogens of important crop species^6^. This subphylum contains the genus *Saccharomyces*, whose members are responsible for baking, brewing, and winemaking industries, which total over a trillion-dollar annual market share. Along with the popular model organism *Saccharomyces cerevisiae*, other emerging yeast models, such as *Komagataella* (*Pichia pastoris), Lipomyces starkeyi, Yarrowia lipolytica,* and *Zygosaccharomyces spp.* are being developed with applications for pharmaceuticals, biofuels, cosmetics, and other biotechnologies^7–11^. 7 of 19 priority fungal pathogens^12^ recently identified by the World Health Organization occur in Saccharomycotina. These include members of the polyphyletic genus *Candida,* which are responsible for over 400,000 life-threatening infections annually with 46-75% mortality^13^.

Despite their relevance to science, technology, industry, and human health, very little is known about the natural distribution of yeast diversity and the factors that govern it^2^. The pathogen *Candida auris* was only described in 2009 but has since been found in 30 countries globally within a decade for unknown reasons^14^. The yeast *Saccharomyces eubayanus*, one of the parental species that gave rise to the lager brewing hybrid *S. pastorianus*, was identified in the wild in 2011^15^, and European populations were only discovered in 2022^16^. Fungi more generally have been traditionally excluded from macroecological studies, and are notably absent from seminal studies on which current theory is based^17–19^. What large scale studies do exist are concentrated in soil fungi, where yeasts accounted for only 0.4% of species^20^. While yeasts can be isolated from soil, their environmental range is far broader and they are commonly found in a variety of substrates and microbiomes across plants, animals, and other fungi^1^. Yeasts have been isolated from locations as diverse as sterile hospital environments^21^ to penguin feces^22^, and can metabolize alcohols, ketones, organic acids, and more^6^. Due to the unique and exceptional diversity of their fundamental niche space, the macroecology of yeasts may differ significantly from other eukaryotic clades. To discover global patterns in yeast diversity and distributions we predicted distribution maps for 186 species and tested drivers across 96 environmental variables.

## Results

To explore the global distribution of yeast species diversity, we used machine learning to infer the full geographic ranges of each species with at least five unique occurrence records. Of these 233 species, 47 with a true positive or true negative rate less than 75% were removed, yielding a total of 186 species representing 9 of 12 Saccharomycotina orders^11^ (Fig. S1). Taxonomic bias is known to confound geographic analyses of species richness^23^. Care must be taken to ensure diversity hotspots are indeed areas of increased species richness, and not just areas of increased taxonomic scrutiny. To assess this possible bias, we compared the observed geographic species richness of the training data defined by the taxonomy used by this study (based on conventional taxonomic standards) to species hypotheses defined by the UNITE database^24^ (based on genetic clustering). Diversity patterns were highly congruent globally between both taxonomies (p<2.2E- 16, r^2^=0.798) (Fig. S2), which indicates that the species richness estimates used by this study reflect true biological patterns. Sampling bias is another factor that can significantly influence biogeographic analyses. The relationship between sampling density was much weaker for predicted diversity estimates (p=6.5E-7, r^2^=0.028) than empirical observations (p=2.2E-16, r^2^=0.402), demonstrating the power of the machine learning approach used by this study to disentangle meaningful phenomena from false signal produced by sampling artifacts (Fig. S3).

Distribution maps predicted through machine learning revealed several distinct hotspots of yeast diversity (Fig. 1), particularly in temperate forests (Fig. 2). Of the 11 most species rich ecoregions all were extratropical forests. Eight were classified as mixed forests and another eight were montane, associated with mountain ranges such as the Alps, Pyrenees, Caucasus, and the Appalachians. Mixed forests harbor the greatest higher-level taxonomic plant diversity, which is thought to contribute heavily to the biodiversity of other fungal groups like ectomycorrhizal mycobionts^25,26^. Similarly, montane ecosystems are known to be exceptionally diverse^27,28^, with radically different assemblages of plants and animals occurring in close proximity along elevational clines.

**Figure 1.**
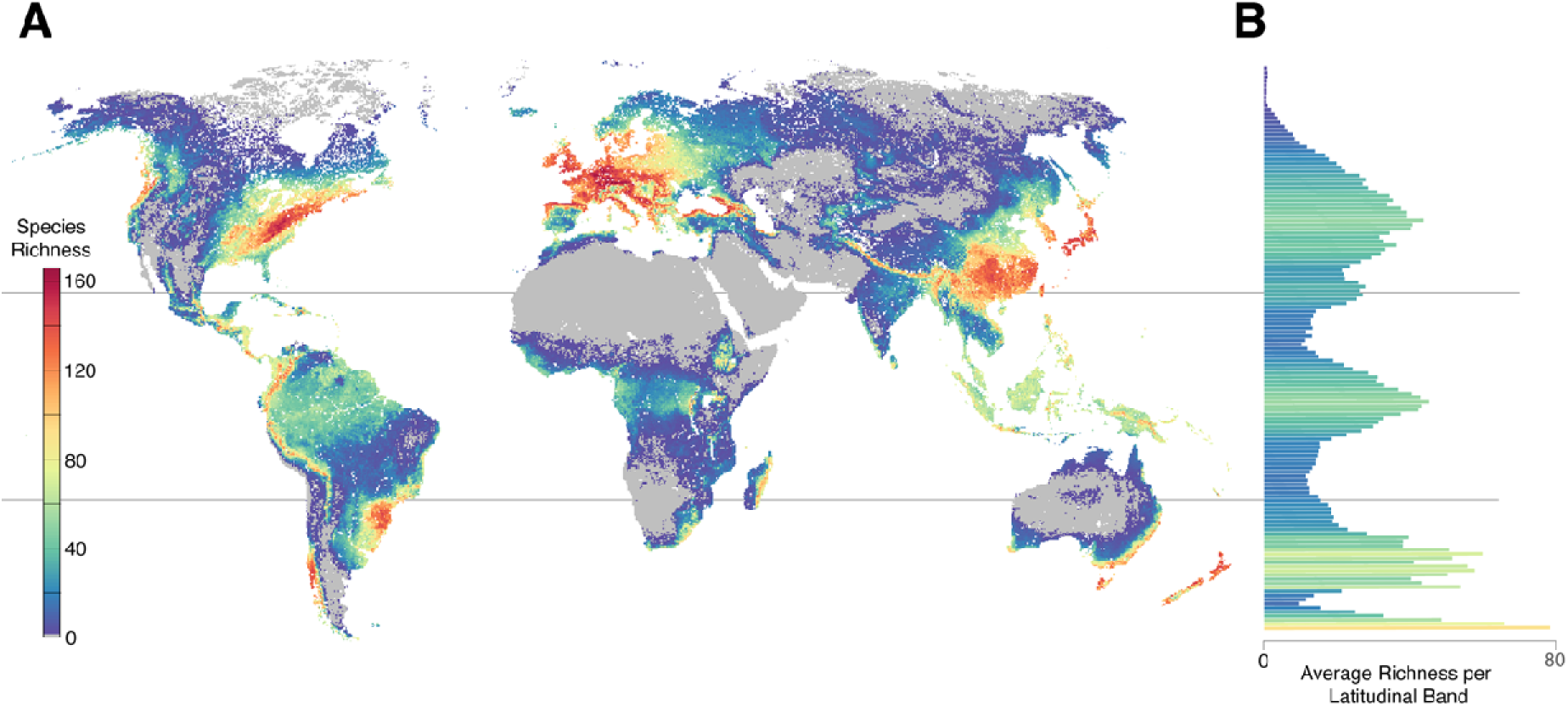
Global yeast diversity. A) Heat map of the distributions of 186 yeast species inferred through random forest machine learning models. B) Average species richness per grid cell for each latitude band or line.

**Figure 2.**
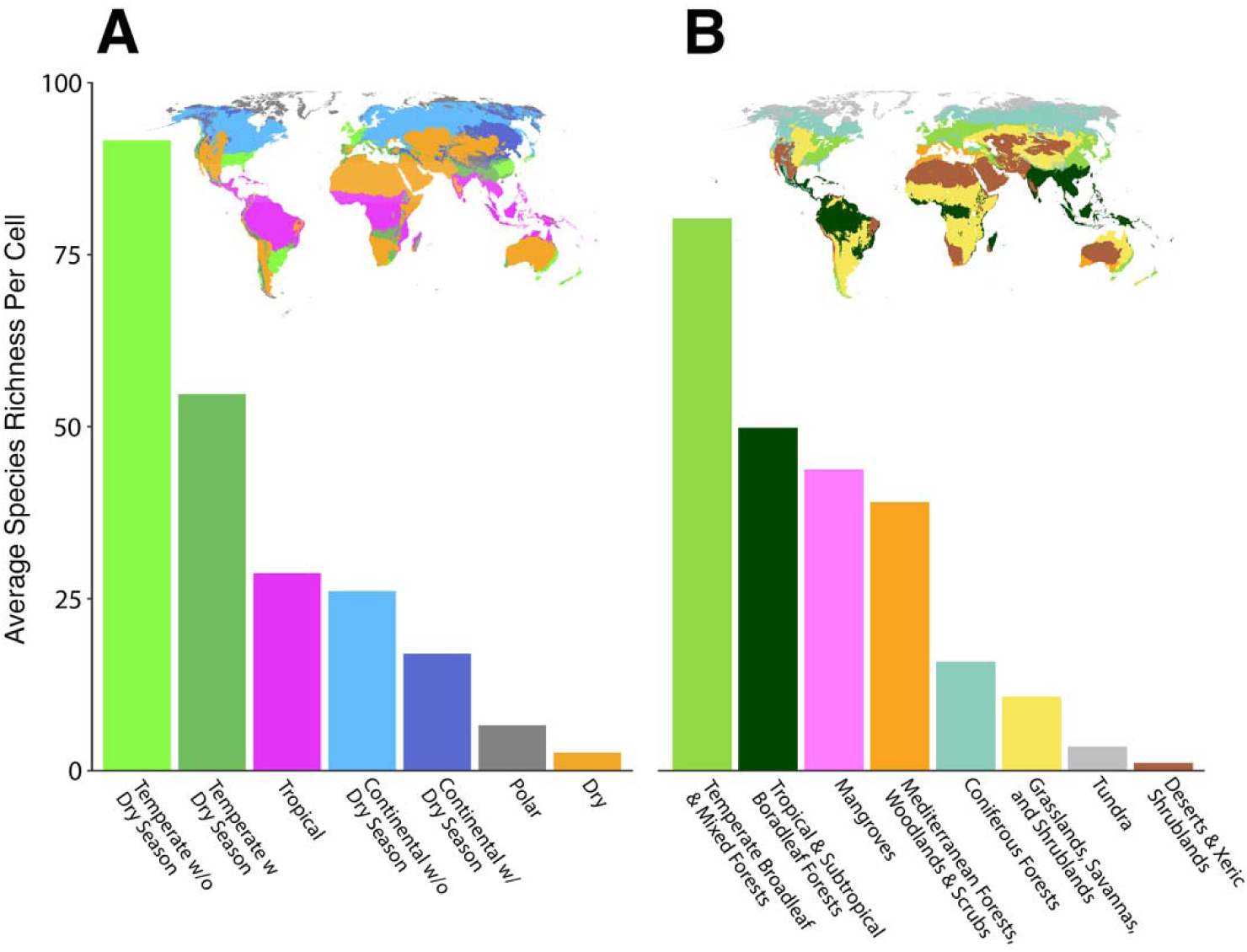
Yeast species richness is concentrated in temperate, mixed forests. Average species richness per grid cell for each Köppen-Geiger climate class (A) and biome (B).

Predicted yeast species richness was highest in mixed, montane forests. To explore which environmental drivers contribute most to yeast diversity in these regions, regression models were performed for 96 variables (Table S1). The heterogeneity in vegetation and topography across montane mixed forests provides a plethora of microhabitats and ecological niches for yeasts to occupy, which may contribute to their high diversity in these environments. This hypothesis is supported by our environmental regression analysis. Two of the variables with 100% relative importance in predicting species richness are enhanced vegetation index diversity and the topography principal component (Fig. 3, Table S2). Plant species richness and geomorphic class diversity were also highly significant, with 98% and 99% relative importance, respectively. By contrast, relative importance for the principal component including forest biomass was just 39%, and altitude was not significant at all (false discovery rate (FDR)=0.33). This result suggests that it is not the forests and mountainous regions per se that are conducive to yeast diversity, but rather the heterogeneity of hosts and landscapes these environments often provide.

**Figure 3.**
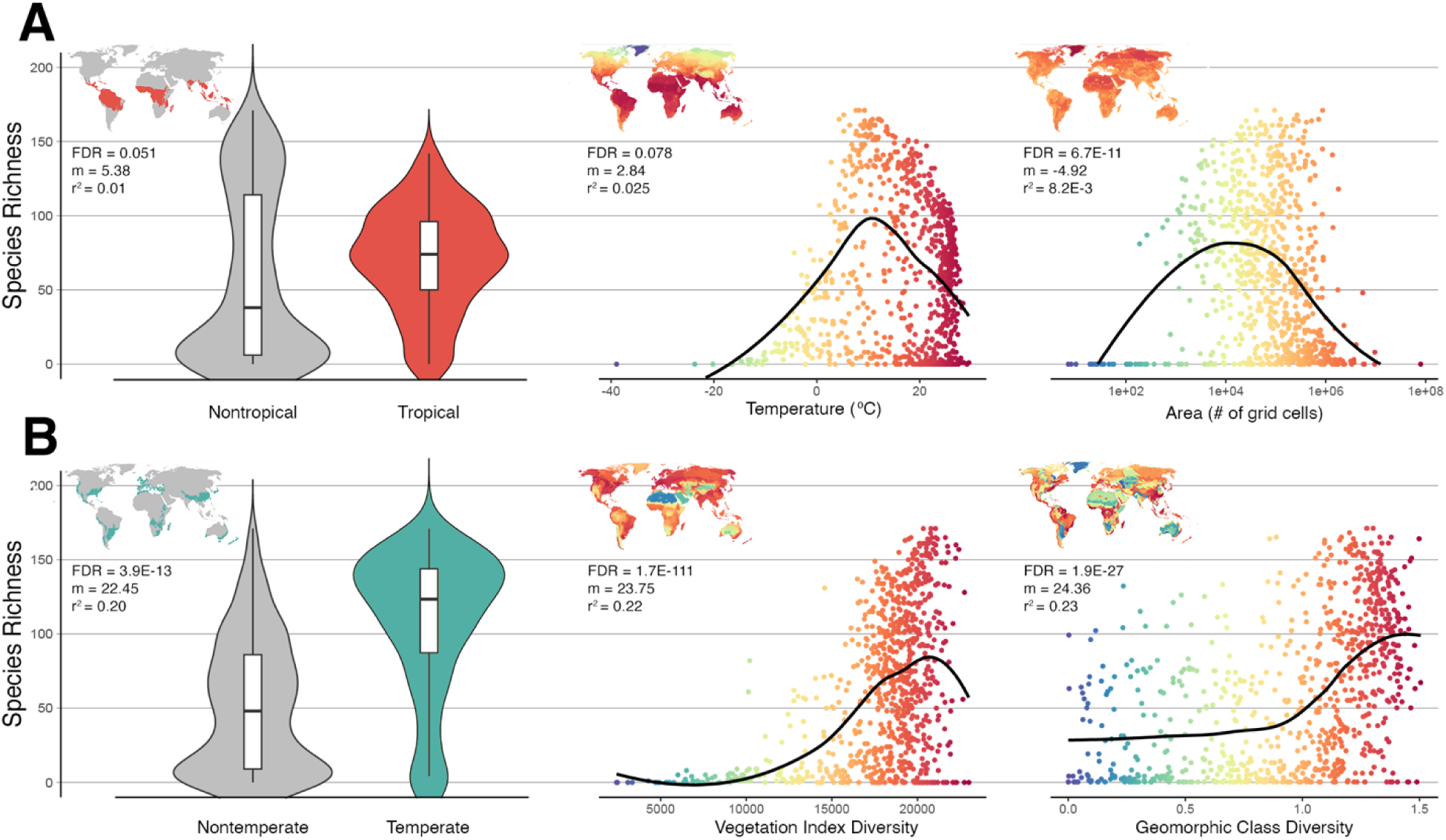
Traditional predictors of species diversity are poor indicators of yeast species diversity. A) Variables that scale with diversity in other clades, such as tropical climates (left), temperature (center), and area (right), did not scale with yeast species diversity. B) Three select variables that were among the best predictors of yeast species diversity: temperate climates (left), vegetation diversity (center), and geomorphic class diversity (right). All graphs represent the same regression analysis with the following summary statistics; FDR: false discovery rate of the negative binomial regression. m: scaled slope of linear regression. r^2^: coefficient of determination for linear regression. Trend-lines were produced through locally weighted smoothing.

Yeast species show extensive variation in their species ranges. For example, *Metschnikowia gruessii* (order Serinales) was predicted to occur in just 9 ecoregions while *Kockiozyma suomiensis* (order Lipomycetales) was predicted to occur in 338, covering over a quarter of the earth’s terrestrial surface (Fig. 4A). We tested three variables that are expected to influence species range size: niche breadth, species richness, and absolute latitude. Niche breadth was obtained from a recent study^4^ that generated experimental growth curves across 18 carbon sources for every species in our dataset. We found that the number of different carbon sources a species was able to metabolize had a significant impact on range size (Fig. 4A). Carbon specialists, which can only grow on a limited number of carbon sources^4^, had significantly (p=0.02) smaller geographic ranges compared to non-specialist species. Range size was also significantly negatively correlated with species richness (p≈0) (Fig. 4B). Species who occupied environments with high numbers of other yeast species were more likely to have smaller ranges. Lastly, absolute latitude had a negative effect (p≈0) on species range size, such that yeast species occupying ecoregions closer to the equator had larger range sizes than more temperate species (Fig. 4C), with species ranges becoming smaller with distance from the equator.

**Figure 4.**
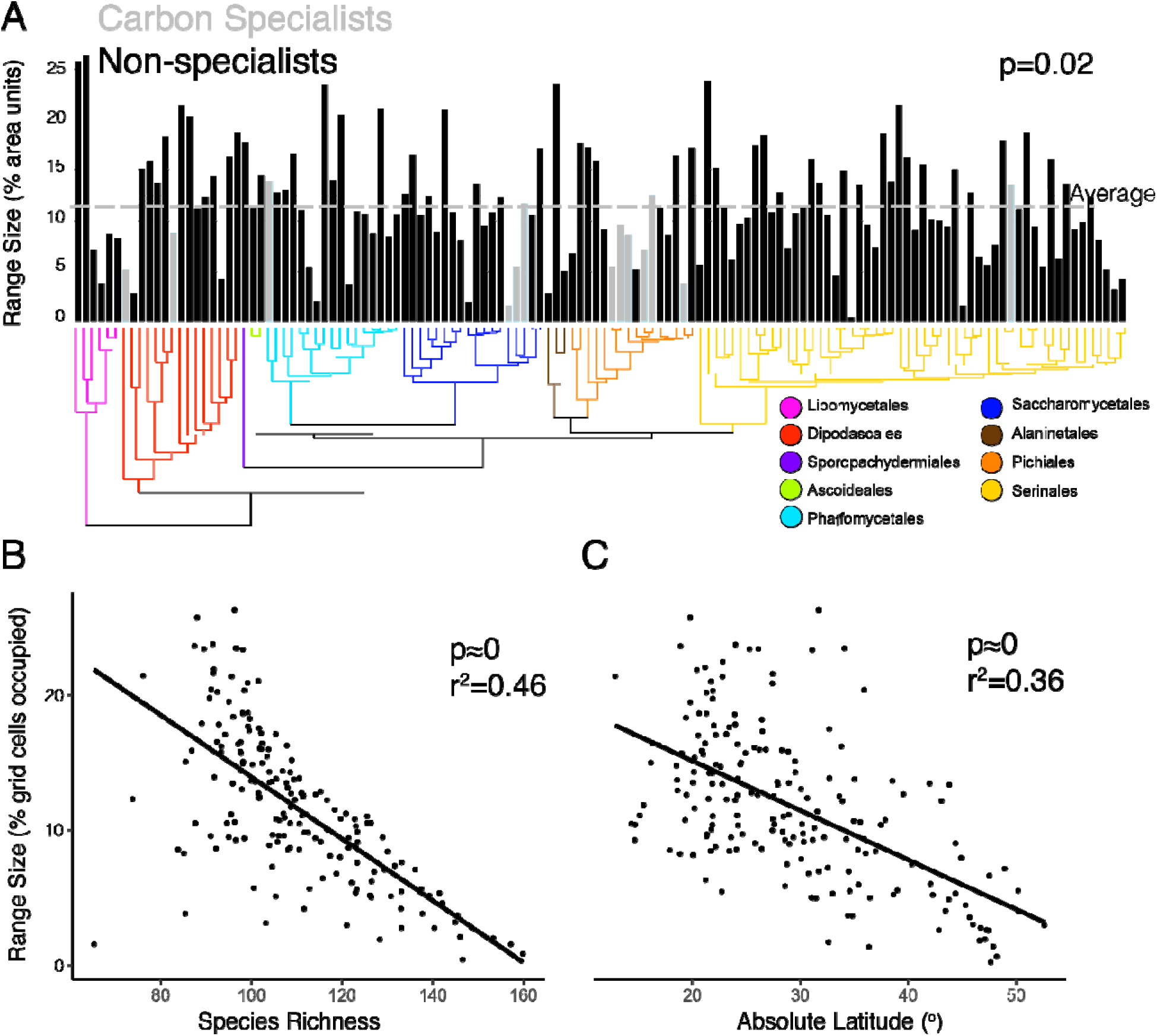
Yeast species range size scales negatively with species richness and latitude, positively with carbon niche breadth. A) Specialist species that grew on only a few carbon sources had significantly smaller geographic ranges than non-specialists. p-values represent a phylogenetic ANOVA test. Size of inferred ranges for each species included in this study, compared to B) species richness, and C) absolute latitude. Summary statistics represented phylogenetic generalized least squares tests. All tests use the same underlying time-calibrated tree from Opulente, LaBella et al. 2023^4^

Yeast species ranges appear to be limited both by high species richness and carbon niche breadth, suggesting that niche partitioning plays an important role in yeast biogeography. Species in high diversity areas have restricted ranges, implying that intraspecific competition limits geographic expansion. The limited range of specialists further demonstrates that the fundamental niche space available to a species has macroecological consequences.

Comparisons of the macroecology of yeasts to other eukaryotic clades reveals several similarities. For example, richness peaks in montane forests^27,28^ and a positive association between niche breadth and range size^17,29^ are general patterns predicted by mainstream macroecological theory. Nevertheless, we also identified three major respects in which yeast macroecology deviates substantially from that of many other eukaryotic groups.

First, it is generally expected that species richness scales with resource availability, usually represented with proxy variables, such as area, temperature, or productivity^30^. However, of these traditional predictors, only productivity emerged as a driver of yeast diversity. Net primary productivity (NPP) had a strong, significant relationship with species richness (FDR=1.1E-57, r^2^=0.24) (Table S2). After highly correlated variables were decomposed into principal components (see Methods), the resulting productivity principal component constructed from net primary productivity, growing season, and soil respiration was similarly predictive, with 100% relative importance (Fig. 3, Table S1).

Surprisingly, neither temperature nor area had a positive effect on yeast species richness. Area size had a significant relationship with richness (FDR=6.8E-11, r^2^=8.2E-3), but the relationship was negative; yeast species richness was greater in smaller ecoregions, which was the opposite trend of what would be expected (Fig. 3). Mean annual temperature had no significant relationship with yeast diversity (FDR=0.078, r^2^=9.4E-3). The temperature-associated principal component constructed from snow cover and energy from the sun was also insignificant (FDR=0.23, r^2^≈0). Temperature was previously identified as an important factor influencing the range of *Saccharomyces* species^31,32^, which has important implications as the ranges of many fungal pathogens are predicted to expand due to climate change^33^. Our analysis suggests that this association between temperature and species range is also true throughout the subphylum since temperature range and temperature mean were the 7^th^ and 9^th^ most important continuous variables in our distribution models, respectively (Table S3). However, while temperature is an important determinant of yeast species distributions, it is not predictive of yeast species diversity globally.

Second, the latitudinal diversity gradient, or the tendency for species richness to peak in tropical climates, is arguably the most widely observed macroecological trend^18^. In Saccharomycotina however, temperate regions held the most diversity with an average species richness of 73.6 species per grid cell, a value 2.6x higher than that of tropical regions (Fig. 2). Additionally, while temperate regions held significantly more richness than non-temperate regions (FDR=3.9E-13, r^2^=0.20), the species richness of tropical regions did not significantly differ from the richness of non-tropical regions (FDR=0.051, r^2^=0.01) (Fig. 3). Previous studies that have observed an inverse latitudinal diversity gradient in other fungal clades have suggested negative relationships between fungal diversity and plant richness^34^ or temperature^35^ as potential drivers. However, as mentioned above, we found that yeast species diversity was positively correlated with plant species richness (FDR=5.9E-43, r^2^=0.22) and uncorrelated with temperature (FDR=0.078, r^2^=1.9E-3).

The absence of tropical diversity in certain fungal clades could also be due to historical biogeographical factors^25^. In ectomycorrhizal fungi, for example, there are no known obligately tropical species^36^, suggesting that lineages originally arose in temperate regions. However, as has been reported in other clades^37,38^, diversity and diversification appear to be only weakly correlated in yeasts, suggesting that historical hotspots of diversification are not necessarily current hotspots of diversity (Fig. S4). Additionally, variables tracking climate changes since the last glacial maximum were largely insignificant and had some of the smallest effect sizes measured. It is possible, that due to the short generation times and widespread dispersal capabilities of many yeast species^39,40^, historical processes that operate over thousands of years have had minimal impact on modern distributions. Such a scenario may also help to explain the absence of a latitudinal diversity gradient. If yeast species can rapidly colonize and saturate environmental niches that were previously unavailable due to climate shifts or glacial cycles, it may explain why species richness is not concentrated in the more stable tropics.

Third, Rapoport’s rule^41^, or the positive relationship between species range size and latitude, was also found to be reversed in yeasts. As mentioned above, distance from the equator had a significant (p≈0), negative relationship with species range size. Though the generality of Rapoport’s rule has been extensively questioned^42,43^, it has been identified as a major factor in the distribution of soil fungi, particularly in Agaricomycetes^20^. Rapoport’s rule was originally postulated in order to explain the latitudinal diversity gradient, since the smaller ranges of species in the tropics would enable more species to coexist. If Rapoport’s rule and latitudinal diversity gradients are indeed connected, it would explain the observed trend of both of them being inverted in yeasts.

In conclusion, we sought to uncover the global diversity and distribution of the Saccharomycotina yeasts. As single-celled organisms, the life history and lifestyle of yeasts are markedly different from many other eukaryotic clades. This divergence is reflected in their macroecology, which sets them apart even from other fungi^20^. We did not find evidence of many commonly observed ecological patterns. Predicted yeast diversity is concentrated in temperate climates, not the tropics. Similarly, species range size decreases with distance from the equator, an inverse of Rapaport’s rule. Additionally, neither temperature, nor area, scale with species richness. These surprising findings emphasize the need in macroecology to study a variety of underexplored clades, especially those with unique life history traits.

The distribution models used by this study are reliant on environmental sampling. Wild yeasts are severely under sampled, which could influence the accuracy of our machine learning predictive models. Nevertheless, our inferences of yeast species richness are consistent with current knowledge. For example, biodiversity hotspots in Western European forests^32^. Perhaps more importantly, our models also make specific predictions that can be tested through additional sampling. Specifically, ecoregions around the Mediterranean and Black seas such as the north Turkish coast and montane forests along the Apennine and Rhodope mountain ranges were in the 98^th^ percentile for yeast species richness, despite having zero samples in our training data. The Appalachian Mountains in the U.S. might similarly be an underappreciated biodiversity hotspot for yeasts, with species richness estimates rivaling that of western Europe despite having less than 3% of their sampling. Guiding both geographic and taxonomic sampling of this important clade toward specific poorly sampled ecoregions will likely greatly increase the resolution and power of future studies. For understudied and under sampled clades like the yeasts, employing a computational predictive framework, such as the one developed in this study, can guide future sampling efforts. We hope that both geographic and taxonomic sampling of this important clade continue to improve, which will help increase the resolution and power of future studies.

While the distribution patterns of yeast diversity are distinct from many other eukaryotes, the threats yeast face may be largely the same. We found that yeast diversity hotspots are characterized by temperate, montane, mixed forests. Notably, these ecosystems are some of the most impacted by human activities and climate change. Forests in central Europe, east Asia, and southwest Brazil, where yeast diversity is high, are dominated by secondary growth^44^, having previously been disturbed by human activities. Similarly, montane environments are particularly impacted by climate change as communities shift upslope in response to rising temperatures, altering species ranges in the process^27,45^. As temperate ecosystems are forced to retreat to higher latitude and altitudes in a warming world, yeast diversity hotspots will need to adapt with them or face extinction. The methodology used by this study is readily adjustable to an array of future climate scenarios, and it may prove useful in assessing how yeast diversity, including economically relevant and pathogenetic species, is affected by past, present, and future anthropogenic transformations.

## Methods

### Dataset

To obtain a comprehensive record of Saccharomycotina biogeographic distribution, several data sources were queried. All available Saccharomycotina occurrence records without flagged geospatial issues were downloaded from the Global Biodiversity Information Facility on December 14, 2022^46^ (doi.org/10.15468/dl.n4fkqs). These data were further filtered by removing any record with a reported coordinate uncertainty of 1km or greater. Saccharomycotina records were also downloaded from the GlobalFungi^47^ dataset (release 4), and two published papers^2,48^. In Spurley et al. (2022), records marked with the “anthropic” flag were removed, as this study is primarily interested in the diversity and distribution of naturally occurring yeasts. Similarly, the industrial hybrid species *Saccharomyces bayanus* and *Saccharomyces pastorianus* were excluded from analysis. Though now considered a naturally-occurring species distinct from *S. bayanus*, *Saccharomyces uvarum* records were also removed as a conservative measure. After records were combined from all four data sources, species names were reconciled with the most recent taxonomy^11^ (Table S4), and two additional filtering steps were applied. First, coordinate resolution needed to be at least two decimal places. Second, the R package CoordinateCleaner was employed to remove suspicious records, such as those with equal latitude and longitude coordinates, zero coordinates, or coordinates matching the centroid of counties/provinces or biodiversity institutions. The full filtered search resulted in 22,355 Saccharomycotina occurrence records, representing every biome on earth and 49.7% of terrestrial ecoregions.

Each occurrence record was associated with 96 environmental variables describing the climate, history, soil, vegetation, and anthropogenic inputs of the region. All variables were taken from publicly available sources and projected onto the WGS84 coordinate system at 30” (∼1 km^2^) resolution. Further details for each variable are available at Table S5. To avoid overfitting or overrepresentation of specific sampling sites in the training data, records with identical environmental variables of the same species within the same hundredth degree of latitude or longitude were aggregated into one. Finally, records with any missing data were also removed, resulting in a training dataset of 12,816 presences.

### Species distribution modeling with machine learning

To infer species occurrences in areas of limited sampling, machine learning random forest models were used. 233 models were constructed, one for every species with at least 5 occurrence records. 100,000 environmental datapoints were randomly sampled as pseudo-absences. Modeling was performed using the R package ‘randomforest’. A down-sampling approach was used for training, which has been shown to reduce overfitting and significantly improve results in species distribution modeling^49^. Each random forest model consisted of 100 decision trees. Otherwise, default parameters were used. A leave-one-out strategy was used for validation, and 186 models with at least a 75% true positive rate and 75% true negative rate were retained for downstream analysis. On average, models for these 186 species had an area under the receiver operating characteristic curve of 0.92, a true positive rate of 87%, and a true negative rate of 90% (Fig. S5). Of the 96 environmental variables used in training, Köppen-Geiger climate classifications were the most predictive, followed by ecofloristic zones, biomes, and soil classifications. Together these four categorical variables represented almost a quarter of all variable importance, with 24.7% of the total mean decrease in Gini index across all variables. We also note that variables that are important for the binary classification task of random forest models are not necessarily those that are the most predictive of overall richness. For example, mean annual temperature was the 9^th^ most important continuous variable for distribution modeling but had an insignificant (FDR=0.078) effect on richness. Conversely, geomorphic class diversity was the 3^rd^ least important continuous variable for distribution modeling but had 100% relative importance to richness regressions.

### Diversity and diversification estimation

To reduce computational costs and to increase interpretability of results, terrestrial ecoregions were selected as the main unit of analysis for this study, which are defined by the World Wildlife Fund as ‘a large unit of land containing a geographically distinct assemblage of species, natural communities, and environmental conditions’^50^. To accomplish this analysis, environmental variables and species richness estimates were aggregated into ecoregions. For the 90 continuous environmental variables in our training dataset, we simply took the mean value of all grid cells in a given ecoregion (Table S6). Select categorical variables were also encoded into 6 binary variables, which were based on the majority class within each ecoregion (Table S7). Species were said to be found in a particular ecoregion if they were predicted to occur in at least 10% of that ecoregion’s grid cells according to the random forest model. Speciation rates were inferred from the DR statistic^51,52^ calculated from the inverse equal splits method^53^, using the time-calibrated phylogeny published in Opulente & LaBella 2023^4^. Ecoregion specific rates were calculated using a weighted mean of speciation rates for all species found in a given ecoregion.

Weights represented the inverse of the number of ecoregions in which a given species occurred, such that species endemic to a specific ecoregion contributed more to that ecoregion’s estimate than a cosmopolitan species^52^.

### Environmental analysis

To determine environmental drivers of yeast diversity, regression models were constructed for each of the 96 quantitative variables, with yeast species richness as the dependent variable in each case. As species richness is always represented by a non-negative integer, negative-binomial regressions were used, which are thought to be more appropriate for count data and, in practice, had consistently better Akaike information criterion scores than linear models. To increase interpretability of summary statistics, scaled linear regressions were also performed, taking r^2^ as a measure for goodness-of-fit and the slope (m) as a measure of effect size. 16 variables whose negative binomial regressions had false discovery rates >0.05 were removed from downstream analysis. To reduce correlations between environmental variables, highly correlated variables were decomposed into single principal components. Effort was made to preserve the interpretation of principal components wherever possible. Each principal component explained at least 83% of the total variance (μ=93%); further details can be found at Table S8. After highly correlated variables were decomposed, the greatest r^2^ between variables was 0.71 (μ=0.11) (Fig. S6). To estimate the contribution of the most predictive environmental variables and principal components, relative importance analysis was used. Negative binomial regression models were constructed from every combination of the 16 variables and principal components whose linear relationship with species richness had r^2^>0.15 and m>0.20; species richness was the dependent variable. This strategy resulted in 65,535 total models. Akaike weights were then calculated and used to estimate relative importance for each predictor^54^.

### Species Range Size Analysis

Several estimates were measured to test drivers of species range size. Species range size itself was estimated as the total fraction of grid cells predicted to be occupied by a given species.

Latitude and species range overlaps were estimated for each species as the average value across every ecoregion in which a given species was predicted to occur (Table S9). Specialist classifications were taken from Opulente, LaBella et al. 2023^4^, which inferred niche-breadth through experimental quantitative growth assays on 18 carbon sources. The positive relationship between niche-breadth and geographic range size has been identified as a major macroecological pattern in plants and animals^17,19^. However, this consensus has also attracted controversy for two main reasons. First, niche-breadth is a broadly defined concept often measured along multiple axes, such as diet, habitat, and tolerance, which are not necessarily correlated^17^. Second, as range size and niche-breadth are typically inferred from the same underlying data (occurrence records), sampling artifacts can produce spurious correlations^29,55,56^. The yeast dataset utilized by this study circumvents both these issues. The external absorption mode of feeding in yeasts^57^ means that diet and habitat are one and the same, providing a convenient and unique lens through which to measure niche-breadth. Additionally, as this study defines niche-breadth independently through experimental growth assays conducted in a laboratory^4^, there is no autocorrelation between niche-breadth and range size. Associations between species range size and diversity/latitude were tested with phylogenetic generalized least squares models implemented in the R package nlme^58^ and niche breadth using phylogenetic ANOVAs implemented in the package geiger^59^.

## Data Availability

All code required to run the species distribution models presented in this paper and replicate primary analyses as well as supplementary data files, including distribution maps and raster files for all 186 species have been deposited online and will be made publicly accessible upon publication.

## Acknowledgements

We thank members of the Rokas Lab and the Y1000+ Project team for helpful discussions throughout the duration of this project. We would also like to thank the Society for the Protection of Underground Networks (SPUN) for their assistance, in particular Dr. Toby Kiers and Dr.

Michael Van Nuland. This work was performed using resources contained within the Advanced Computing Center for research and Education at Vanderbilt University in Nashville, TN.

