## Supplementary Figures and Supplementary Table Legends for "Saccharomycotina yeasts defy longstanding macroecological patterns"

### Supplemental Figures

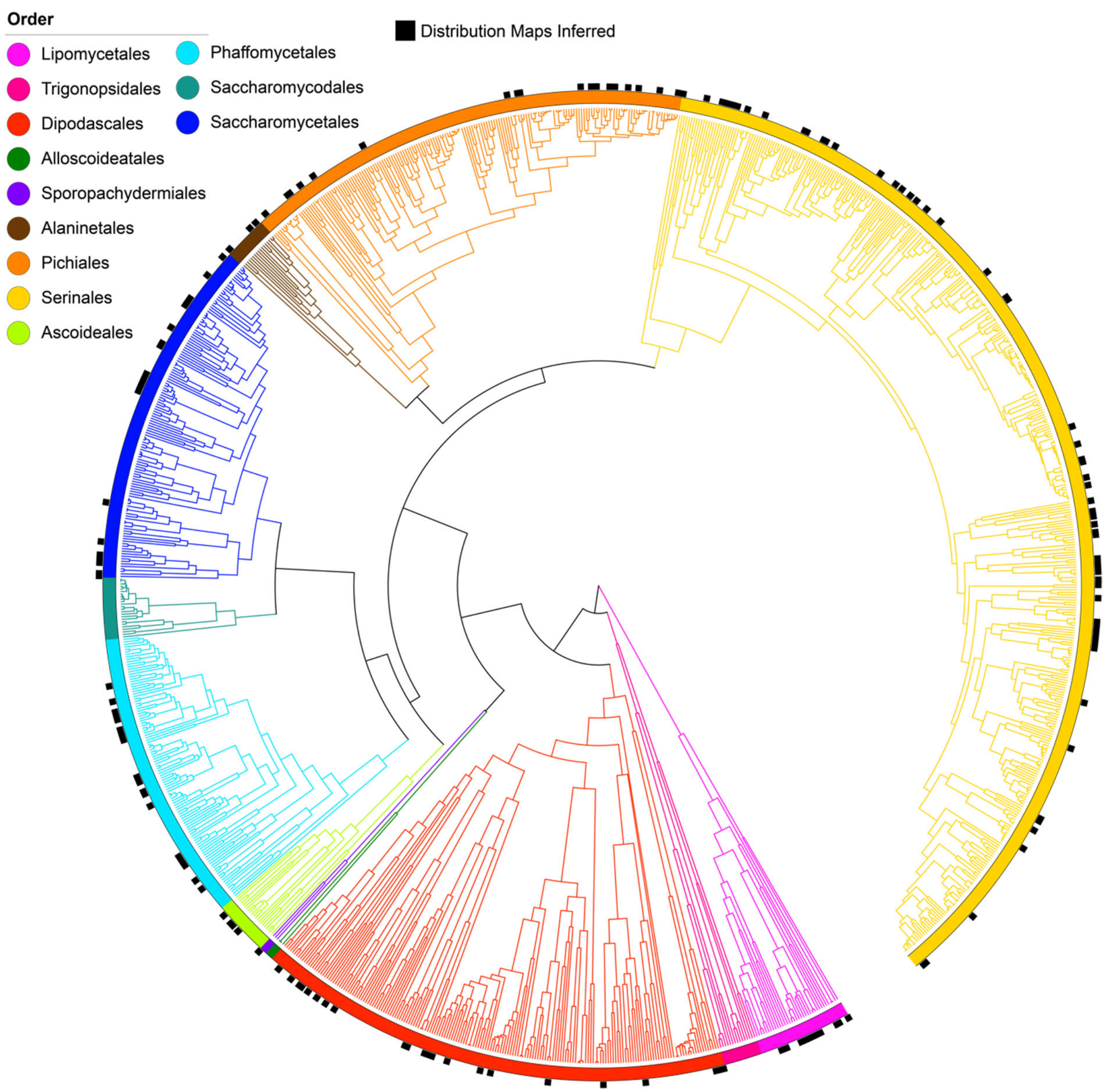


**Figure S1. Phylogenetic distribution of sampled species.** Species with distribution maps inferred by this study mapped onto a phylogeny highlighting the 12 orders of Saccharomycotina. 11 additional species without phylogenetic information were also included.

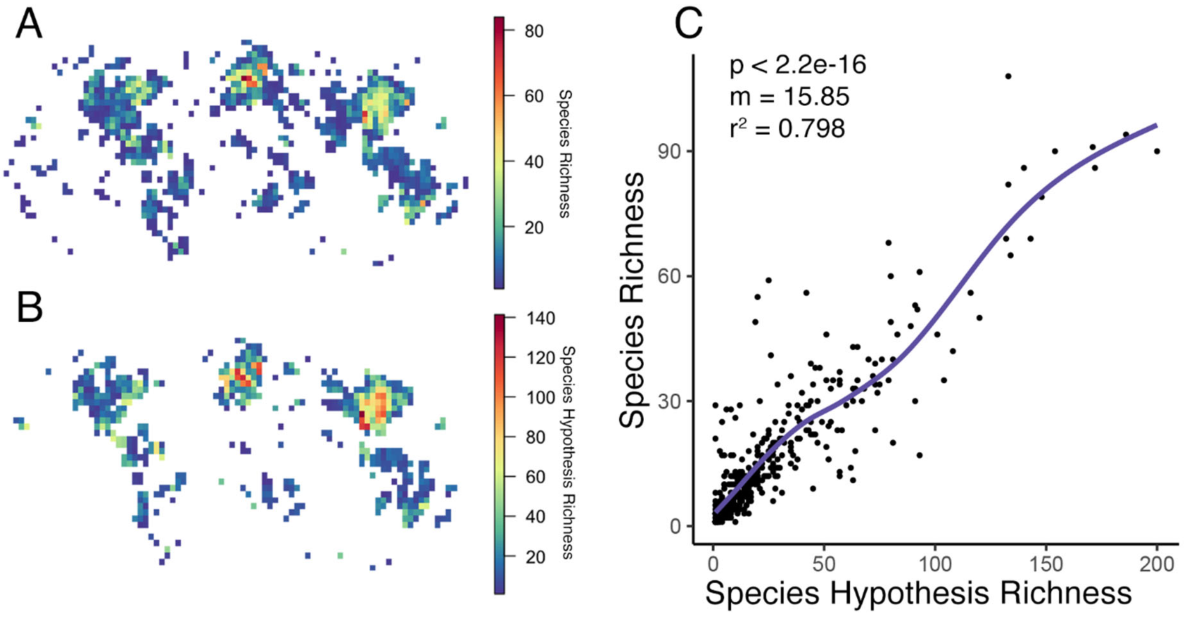


**Figure S2. Global patterns of traditional taxonomy and species hypotheses are congruent.** A) Geographic heat map of observed species richness for all species considered by this study. B) Geographic heat map of species hypotheses as defined by the UNITE database for molecular identification of fungi. C) Correlation between diversity as estimated by species richness and species hypothesis richness. The p-value, scaled slope, and correlation coefficient of the linear model are displayed.

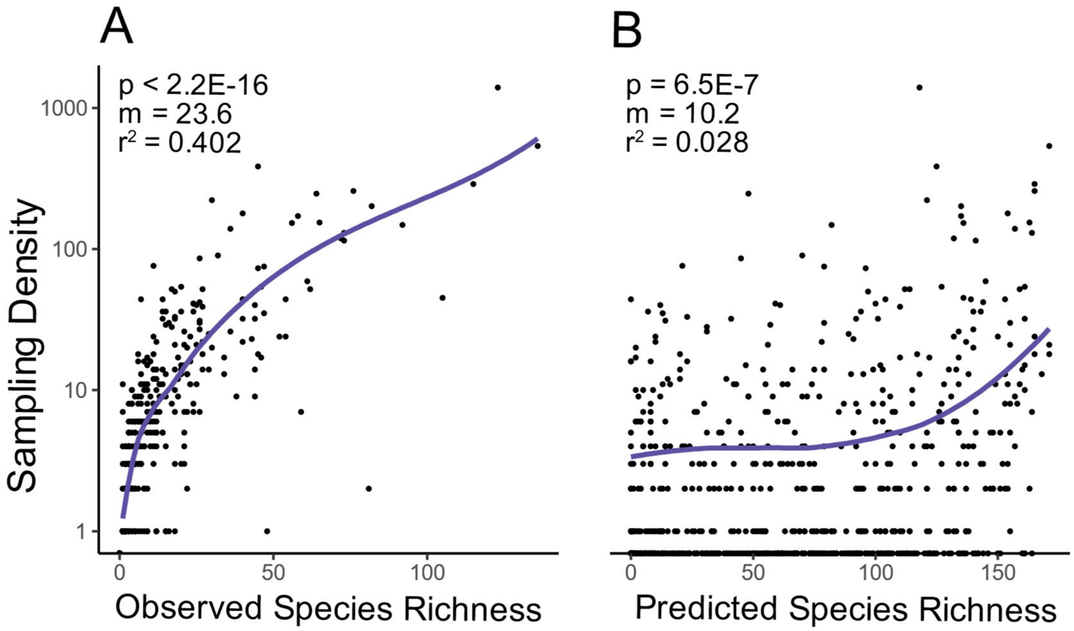


**Figure S3. Machine learning reduces sampling bias**. Sampling effort of each ecoregion versus A) the number of observed species in the training data and B) the number of predicted species as inferred by random forest species distribution models. The p-value, scaled slope, and correlation coefficient of the linear model are also displayed.

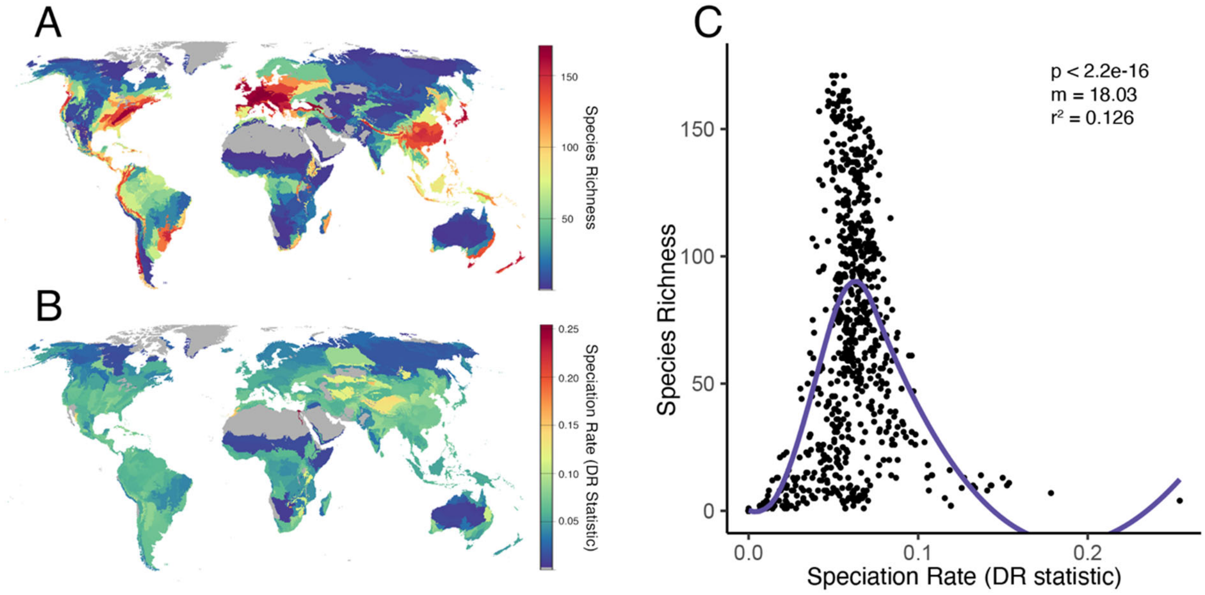


**Figure S4. Diversity and diversification are not strongly correlated.** A) Species richness per ecoregion. B) Speciation rate per ecoregion, as estimated by the DR species-specific test statistic, weighted by species range. C) Correlation between species richness and speciation rate. The p-value, scaled slope, and correlation coefficient of the linear model are displayed.

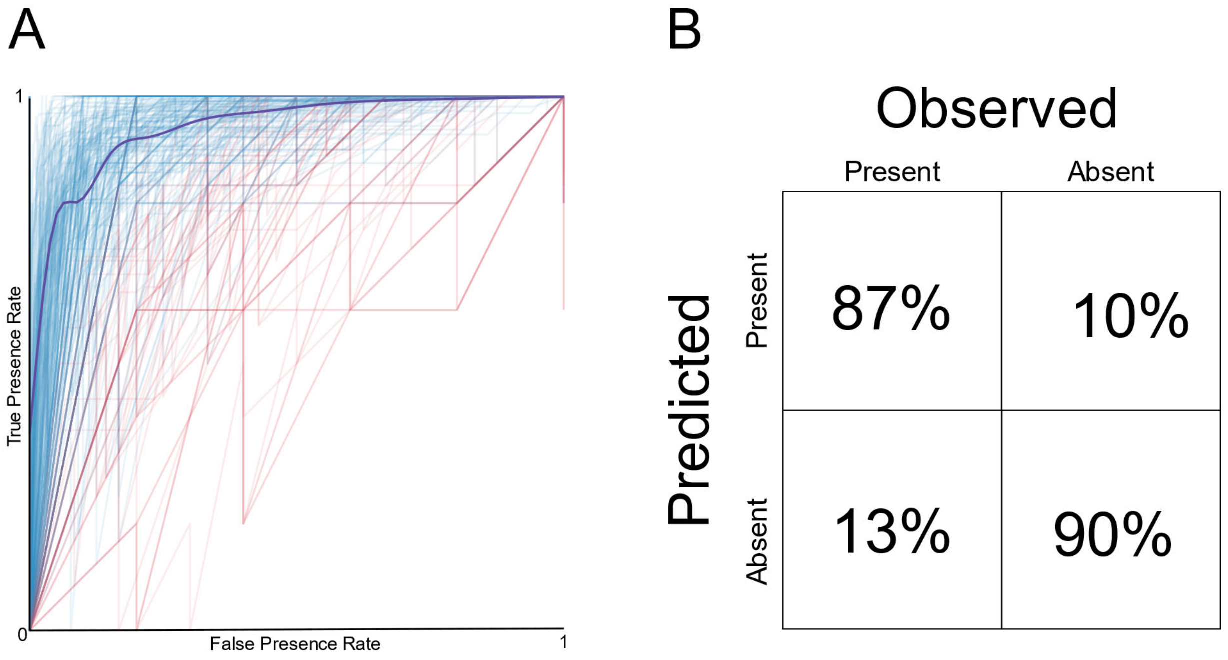


**Figure S5. Machine learning results.** A) Receiver operating characteristic curves of each species considered; species in red fell below our threshold and were excluded from analysis. B) Confusion matrix averaged across all species included in the analysis.

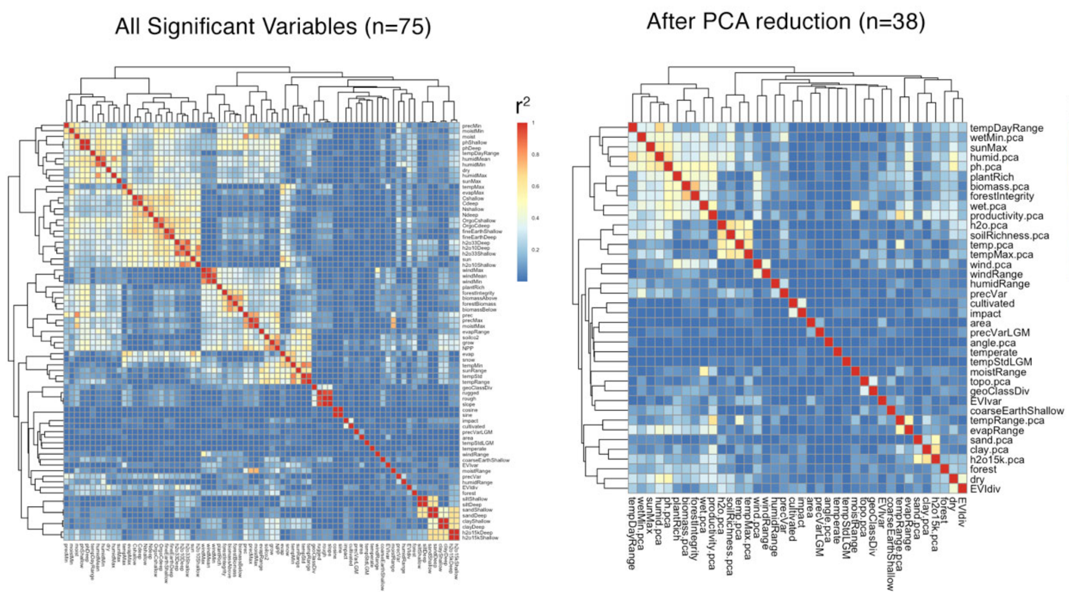


**Figure S6. PCA decomposition reduces correlation between environmental predictors.** Correlation matrix between each significant environmental variable before and after PCA decomposition.

### Supplemental Table Legends

**Table S1. Summary statistics of diversity regression analysis for all variables.** var=predictor name (see Table S5), p=p-value of negative binomial regression, FDR=false discovery rate of negative binomial regression, m=slope of scaled linear regression, r2=coefficient of determination for scaled linear regression.

**Table S2. Summary statistics of diversity regression analysis for significant variables and principal components.** var=predictor name (see Tables S5 and S8), p=p-value of negative binomial regression, FDR=false discovery rate of negative binomial regression, m=slope of scaled linear regression, r2=coefficient of determination for scaled linear regression, relative importance=relative importance of top performing predictors.

**Table S3. Relative importance of training variables.** Mean decrease in Gini index of every training variable for the 186 species distribution models performed by this study.

**Table S4. Species synonyms.** Updated taxonomy used to reconcile previously published occurrence records.

**Table S5. Variable details.** Definitions and details for each environmental variable used in training and diversity regression analysis.

**Table S6. Ecoregion data.** Aggregated environmental variables for each ecoregion, this data underlies the regression analysis (Tables S1 and S2).

**Table S7. Binary variable definitions.** Definitions for select categorical variables encoded as binomial.

**Table S8. Principal component definitions.** Definitions and details for constructed principal components of highly correlated variables.

**Table S9. Range size data.** Range size, mean latitude, and mean species overlap for each species, this data underlies the range size phylogenetic comparative modeling analysis (Fig 4).
